# Sensitive visualization of SARS-CoV-2 RNA with CoronaFISH

**DOI:** 10.1101/2021.02.04.429604

**Authors:** Elena Rensen, Stefano Pietropaoli, Florian Mueller, Christian Weber, Sylvie Souquere, Pierre Isnard, Marion Rabant, Jean-Baptiste Gibier, Etienne Simon-Loriere, Marie-Anne Rameix-Welti, Gérard Pierron, Giovanna Barba-Spaeth, Christophe Zimmer

## Abstract

The current COVID-19 pandemic is caused by the severe acute respiratory syndrome coronavirus 2 (SARS-CoV-2). The positive-sense single-stranded RNA virus contains a single linear RNA segment that serves as a template for transcription and replication, leading to the synthesis of positive and negative-stranded viral RNA (vRNA) in infected cells. Tools to visualize viral RNA directly in infected cells are critical to analyze its replication cycle, screen for therapeutic molecules or study infections in human tissue. Here, we report the design, validation and initial application of fluorescence *in situ* hybridization (FISH) probes to visualize positive or negative RNA of SARS-CoV-2 (CoronaFISH). We demonstrate sensitive visualization of vRNA in African green monkey and several human cell lines, in patient samples and human tissue. We further demonstrate the adaptation of CoronaFISH probes to electron microscopy (EM). We provide all required oligonucleotide sequences, source code to design the probes, and a detailed protocol. We hope that CoronaFISH will complement existing techniques for research on SARS-CoV-2 biology and COVID-19 pathophysiology, drug screening and diagnostics.

## Introduction

Coronavirus disease (COVID-19) emerged by the end of 2019 in Wuhan, China, and led to more than 100 million infections and over 2.1 million deaths (Johns Hopkins University Dashboard). Its causative agent, Severe acute respiratory syndrome coronavirus 2 (SARS-CoV-2), is an enveloped, positive-sense, single-stranded RNA virus. Upon infection, viral replication occurs in the host cell’s cytoplasm, which is massively reorganized (V’kovski *et al*, 2020). The genomic positive-strand viral RNA serves as a template for transcription and replication. The virus synthesizes its own RNA-dependent RNA polymerase (RdRP) to generate negative-sense RNA replication intermediates. This negative strand acts as template for replication of new full-length positive-stranded RNA genomes and for transcription of several smaller, subgenomic positive-stranded RNAs (sgRNAs). These sgRNAs are then used to synthesize all other viral proteins in spatially confined replication complexes. Mature virions are exocytosed and released from the infected host cell. Despite recent progress, many aspects of the SARS-CoV-2 viral replication cycle, including the subcellular location of viral RNA synthesis, are still not fully understood and under active investigation (V’kovski *et al*, 2020).

Several established techniques allow studying SARS-CoV-2 and its interaction with its host. Immunofluorescence (IF) permits the visualization of viral and host proteins in the spatial context of a single cell. However, the development of specific antibodies against novel viruses is time- and cost-intensive, especially if specificity over other closely related viruses is required. Further, the presence in cells of structural viral proteins, such as the Spike protein, does not necessarily imply active viral replication (Cheung *et al*, 2005; Tang *et al*, 2020) and their subcellular localization may not reflect that of the vRNA strands. Other molecular methods, such as RT-PCR, provide an accurate, quantitative readout of viral load and replication dynamics, but are bulk measurements over large cell populations that mask variability between cells and provide no information about the subcellular localization of the virus. RNAseq permits a complete view of the transcriptome of both the host and virus, including in single cells, albeit again without spatial information (Kim *et al*, 2020; Bost *et al*, 2020).

Unlike immunostaining, PCR or sequencing methods, fluorescence *in situ* hybridization (FISH) offers the capacity to directly and specifically visualize viral RNA in single cells (Raj *et al*, 2008; Itzkovitz & van Oudenaarden, 2011; Mueller *et al*, 2013). In RNA-FISH, single RNA molecules are typically targeted with 10–50 fluorescently labeled probes consisting of short (20-30 nucleotides), custom synthesized oligonucleotides with bioinformatically designed sequences. Individual RNAs are subsequently visible as bright, diffraction-limited spots under a microscope, and can be detected with appropriate image analysis methods (Raj *et al*, 2008; Mueller *et al*, 2013). We recently introduced smiFISH (Tsanov *et al*, 2016), an inexpensive variant of this approach that has been used in biological samples ranging from single-cell organisms such as bacteria and yeast to whole tissue sections and organs (Trcek *et al*, 2012; Wu *et al*, 2018; Wang, 2019; Chen *et al*, 2016; Skinner *et al*, 2013), and is ideally suited to visualize RNA viruses and study their subcellular localization and kinetics in host cells (King *et al*, 2018; Rensen *et al*, 2020).

In this study, we designed and validated smiFISH probes against the positive and negative RNA strands of SARS-CoV-2 (CoronaFISH). We demonstrate highly specific viral detection in cell culture, in patient isolates, and in tissue samples. We further demonstrate the flexibility of these probes by adapting them for electron microscopy *in situ* hybridization (EM-ISH). CoronaFISH provides a flexible, cost-efficient and versatile platform for studying SARS-CoV-2 replication at the level of single cells in culture or in tissue, and can potentially be employed for drug screening and diagnosis.

## Results

### Design of probes specific for SARS-CoV-2

Our RNA-FISH approach employs two types of bioinformatically designed DNA oligonucleotides (oligos) (Tsanov *et al*, 2016): (i) unlabeled primary oligos consisting of two parts: a specific sequence complementary to a selected subregion of the target RNA and a readout sequence that is identical among all primary oligos (FLAP sequence), (ii) a fluorescently labeled secondary oligo complementary to the FLAP sequence, allowing visualization by light microscopy. These oligos are hybridized to each other *in vitro* before their use for cellular imaging (**Fig 1a**).

**Figure 1.**
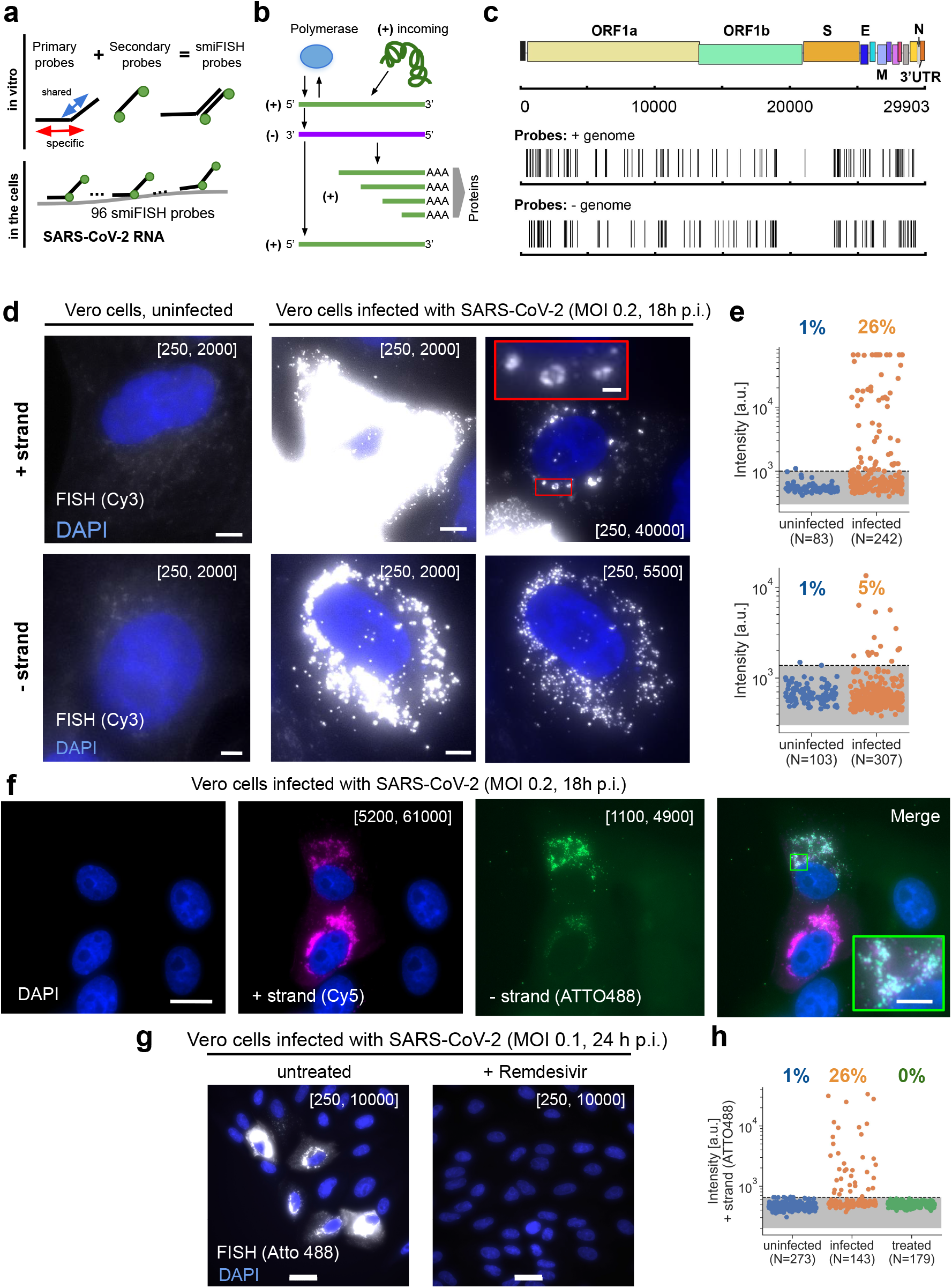
Visualizing SARS-CoV-2 with CoronaFISH. **(a)** Principle of CoronaFISH. 96 primary probes are pre-hybridized *in vitro* with dye-carrying secondary probes via the FLAP sequence. Resulting duplexes are subsequently hybridized in cells to target the SARS-CoV-2 positive or negative RNA. **(b)** Replication cycle of SARS-CoV-2. Incoming, genomic positive-strand RNA is used to produce viral polymerase. Polymerase produces a negative-strand replication intermediate, which serves as a template for synthesis of full-length positive-strand and shorter sub-genomic RNAs. The latter are used to produce other viral proteins. **(c)** Genome of SARS-CoV-2 with indicated probe positions targeting the positive and negative strand. **(d)** Images of uninfected and infected Vero cells with either the positive or negative strands detected with Cy3-labeled probes. Shown are zoom-ins on individual cells. Full-size images in **Fig EV1a**. First column shows uninfected control, second and third column infected cells with different intensity scalings as indicated in brackets (the first and second values in brackets indicate the pixel values corresponding to the lowest and brightest intensities in the displayed image, respectively). Scale bars 5 μm. Scale bar in red inset 1 μm. **(e)** Quantification of signal intensities in individual cells. Dashed line is the 99% quantile estimated from uninfected samples. **(f)** Simultaneous imaging of positive and negative strands with dual-color CoronaFISH. Scale bar in full image 10 μm, in inset 2 μm. **(g)** Images of Remdesivir treated (right) or untreated cells (left). Scale bars 30 μm. (**h**) Quantification of Remdesivir treated cells performed as in e).

A cell infected by SARS-CoV-2 can contain the incoming positive strand, the negative-strand replication intermediate, as well as replicated full-length and sub-genomic positive-strand RNA molecules (**Fig 1b**). We designed two sets of 96 probes, one against the positive strand, and one against the negative strand of SARS-CoV-2 (**Fig 1c**). For more details on the probe design workflow, see the Methods section and the source code (https://github.com/muellerflorian/corona-fish). In brief, we identified an initial list of more than 600 potential probe sequences with our previously described method Oligostan (Tsanov *et al*, 2016). We then further screened these probes to be robust to known genomic variations of SARS-CoV-2 (as of April 2020), while removing probes with affinity to other known β-coronaviruses or viruses frequently causing similar respiratory diseases in human such as Influenza. Lastly, we removed probes overlapping with the transcriptome of several relevant host organisms (human, mouse, African green monkey, hamster and ferret). To establish the final list of 96 probes (**Fig 1c**), we chose probes targeting regions with the highest NGS coverage. The complete list of probe sequences is provided in **Supplementary Table S1**.

### Visualization of SARS-CoV-2 in Vero cells with CoronaFISH

To test our probes, we first used Vero cells (African green monkey), which are known to be permissive for SARS-CoV-2 replication (Takayama, 2020) (see Methods). We processed the samples for FISH following the protocol detailed in **Supplementary Note 1** and imaged cells 18h post infection (p.i.) with a multiplicity of infection (MOI) of 0.2, as well as uninfected control cells. The positive-strand and negative-strand probe-sets were both labeled with the fluorophore Cy3 and imaged separately in distinct experiments.

In uninfected samples, cells displayed only weak and diffuse fluorescent signal when labeled with either probe-set, consistent with unspecific background labeling common in RNA-FISH (Tsanov *et al*, 2016), and occasional bright spots could be observed, mostly located outside cells, presumably due to unspecific probe aggregation (**Fig 1d-e** and **S1a**). By contrast, in infected samples, a large proportion of cells showed very strong and localized signal in large regions of the cytoplasm (**Fig 1d-e** and **S1a**). Quantification of the fluorescent intensities per cell (see Methods) indicated that 26% of infected cells labeled for the positive strand (n=242) had intensities exceeding a threshold that excluded >99% of uninfected cells (n=83) (**Fig 1e**). The fluorescence intensity in these cells was on average 23 times higher than this threshold (s.d. 27) (**Fig 1e**). For the negative strands, we counted 5% of Vero cells with intensities above the similarly defined threshold in infected samples (n=307), vs. <1% for uninfected cells (n=103), with on average 2.5 times higher intensities (s.d. 2.2).

The fluorescent signal for the positive-strand was remarkably strong compared to the uninfected control images (**Fig 1e**). This is consistent with an extremely high per-cell viral yield, which has previously been reported for Vero cells (Ogando *et al*, 2020). The RNA signal was perinuclear and restricted to the cytoplasm, consistent with cytoplasmic replication, as previously reported for other *Coronaviridae* and recently for SARS-CoV-2 (Snijder *et al*, 2020; Fung & Liu, 2019; Stertz *et al*, 2007; Klein *et al*, 2020). Interestingly, we observed bright foci of different sizes and intensities, some of which displayed hollow structures reminiscent of the replication organelles (RO) or double-membrane vesicle (DMV) structures described previously (Ogando *et al*, 2020; Klein *et al*, 2020). The signal of the negative strand was substantially dimmer than for the positive-strand (**Fig 1e**), in agreement with previous reports that the replication intermediate negative strand is less abundant (Wolff *et al*, 2020), but potentially also reflecting diminished labeling efficiency of this probe set. The negative-strand RNA likewise forms clusters of different sizes and intensities in the proximity of the nucleus (**Fig 1d, Fig EV1a**).

We next wanted to demonstrate the ability of CoronaFISH to simultaneously visualize two RNA species with dual-color imaging (Tsanov *et al*, 2016). For this purpose, we used different FLAP sequences on the primary probes set (FLAP-X for positive-strand, and FLAP-Y for negative-strand probes), and labeled them with spectrally different fluophores (Atto488 and Cy5). We then imaged infected Vero cells at 18 h p.i. These images clearly show the presence of both positive and negative RNA strands in the same cells and the same subcellular regions, although the relative abundance of both strands appeared to vary from cell to cell and colocalization was only partial (**Fig 1f, Fig EV1b**). The ability to visualize positive and negative RNA together opens the door to analyzing the intracellular organization of viral transcription and replication dynamics.

Our data thus show that CoronaFISH probes can sensitively and specifically detect the positive and negative strands of SARS-CoV-2 in infected Vero cells.

### Applicability of CoronaFISH to drug screening

We next asked if CoronaFISH can be used to test pharmacological treatments against SARS-CoV-2. To this end, we treated SARS-CoV-2 infected Vero cells with Remdesivir, an adenosine nucleoside triphosphate analog that reduces viral replication *in vitro* by inhibiting the RdRP (Gordon *et al*, 2020). We first established an inhibition curve that showed, in our system, an IC50 concentration of 2.8 µM for Remdesivir (**Fig EV1c**). We then performed FISH against the positive strand using Atto-488 labeled probes in infected cells left untreated or treated with 10 µM of Remdesivir. In untreated samples, 26% of cells (n=143) displayed a strong fluorescence signal (above the 99% threshold defined for n= 273 uninfected cells), as above, whereas Remdesivir-treated cells only showed background signal similar to uninfected cells (0% of n=179 cells above threshold) (**Fig 1g**,**e**). These data suggest that our CoronaFISH probes can be used to test molecules for their ability to inhibit SARS-CoV-2 replication.

### Detection of SARS-CoV-2 RNA in human cell lines

We next tested the probes in several human cell lines known to be permissive to SARS-CoV-2 (Takayama, 2020): Caco-2 (human intestinal epithelial cells), Huh7 (hepatocyte-derived carcinoma cells), and Calu-3 (human lung cancer cells). Each cell line was infected with MOI 0.2 and fixed at 36 h p.i. Titration on the supernatants of the infected cells revealed vastly different replication efficiencies as tested by focus forming assay (**Table 1**). Caco2 and Huh7 cells showed rather low virus levels in the same order of magnitude as Vero cells (2-5×10^3^ FFU/ml), while Calu3 showed two orders of magnitude higher levels of viral RNA (2×10^5^ FFU/ml).

**Table 1.** Titration on the supernatant of infected mammalian cells.

| Cell line | Viral titer [FFU/ml] |
| --- | --- |
| Caco2 | $2 \times 10^3$ |
| Huh7 | $4 \times 10^3$ |
| Calu3 | $2 \times 10^5$ |
| Vero | $5 \times 10^3$ |

We then performed FISH against the positive RNA strand. As for the Vero cells above, the uninfected controls of all human cell types showed only a weak background signal, while a strong signal could be detected in the infected cells (**Fig 2a-c**). However, depending on the cell-type, the number of infected cells, as well as the amount and cellular localization of vRNA detected by CoronaFISH were very different, consistent with different replication dynamics of SARS-CoV-2 in these cell lines. In Caco-2 cells, only a minority of cells appeared infected (19% of n=1,752 cells above intensity threshold defined from n=229 uninfected cells as above), displaying rather low-intensity values, indicating a low abundance of positive-strand viral RNA (6.2-fold above threshold, s.d. 6.8). Huh7 cells were more permissive for viral infection manifesting in more cells displaying vRNA (74% of n=546 above threshold defined on n=246 uninfected cells) and higher RNA signal intensity (11-fold above threshold, s.d. 10). Lastly, all Calu cells were infected (100% of n=773 cells above threshold defined on n=479 uninfected cells) and the signal intensity was higher than in Caco-2 and Huh7 cells (28-fold above threshold, s.d. 14). These data show that CoronaFISH probes can also be used in cell lines of human origin with similar detection performance as in Vero cells.

**Figure 2.**
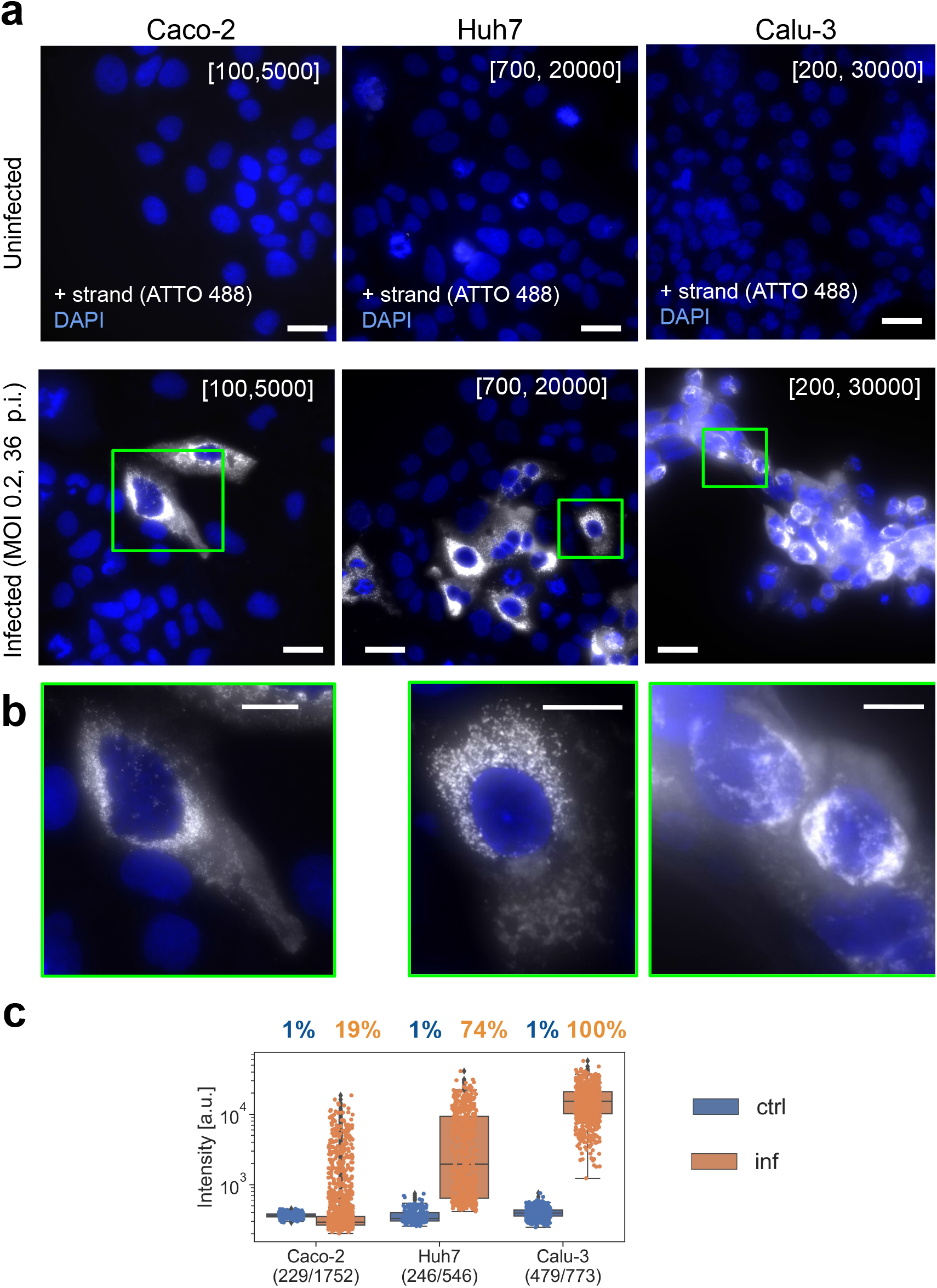
CoronaFISH in human cell lines. **(a-b)** FISH against the positive strand of SARS-CoV-2 in three different cell lines. (**a**) Full field of views, scale bars 30 μm, (b) zoom-ins on indicated green rectangles, scale bars 10 μm. **(c)** FISH signal intensities of cells in a small region around the nucleus of each cell. Box plots as in **Fig 1e**.

### Detection of SARS-CoV-2 RNA in human tissue

Next, we aimed to test if our approach can be used to detect SARS-CoV-2 RNA in samples from patients. The major histopathological finding of the pulmonary system of post mortem COVID-19 patients with acute respiratory distress syndrome (ARDS) is diffuse alveolar damage in the acute or organizing phases. Lung tissue examination mainly shows evidence of intra-alveolar hemorrhage and edema with fibrin and hyaline membranes developed on alveolar walls at the acute phase and proliferative and fibrotic lesions in the alveolar septal walls at the organizing phase (Hanley *et al*, 2020; Bradley *et al*, 2020). However, these lesions are common to multiple forms of ARDS and not specific to COVID-19 and do not shed light on the underlying etiology. CoronaFISH therefore has the potential to specifically detect SARS-CoV-2 infection and characterize viral tropism within the tissue.

As a negative control, we imaged a tissue section sample obtained from a deceased adult patient with diffuse alveolar damage from ARDS prior to the COVID-19 pandemic (see Methods). Histological analysis showed diffuse alveolar damage with an important alveolar hemorrhage, an intra-alveolar and interstitial edema with polymorphic inflammatory infiltrate and the presence of early hyaline membrane adjacent to alveolar walls (**Fig EV2a**). When staining this sample with the positive-strand CoronaFISH probes, no strong fluorescent signal was detected, despite the presence of alveolar damage comparable to patients suffering from COVID-19 (**Fig EV2 b-c**).

As a positive control, we used samples from a COVID-19 patient who died 3 days after admission to the intensive care unit. Histological analysis showed diffuse alveolar damage at the organizing phase with intra-alveolar hyaline membranes and fibrin together with interstitial fibrotic lesions with polymorphic inflammatory cell infiltrate of alveolar walls (**Fig EV2d**). Whereas histology by itself is not sufficient to diagnose lung tissue infection, CoronaFISH revealed infected cells with large cytoplasmic RNA aggregates (**Fig 3a**), illustrating that viral presence can also be detected in tissue sections. Because of the extensive destruction of tissue architecture, determining the affected area of the parenchyma is challenging. However, the cell types (e.g. macrophages or type 2 pneumocytes) infected by SARS-CoV-2 could be revealed by combining CoronaFISH with immunostaining.

**Figure 3.**
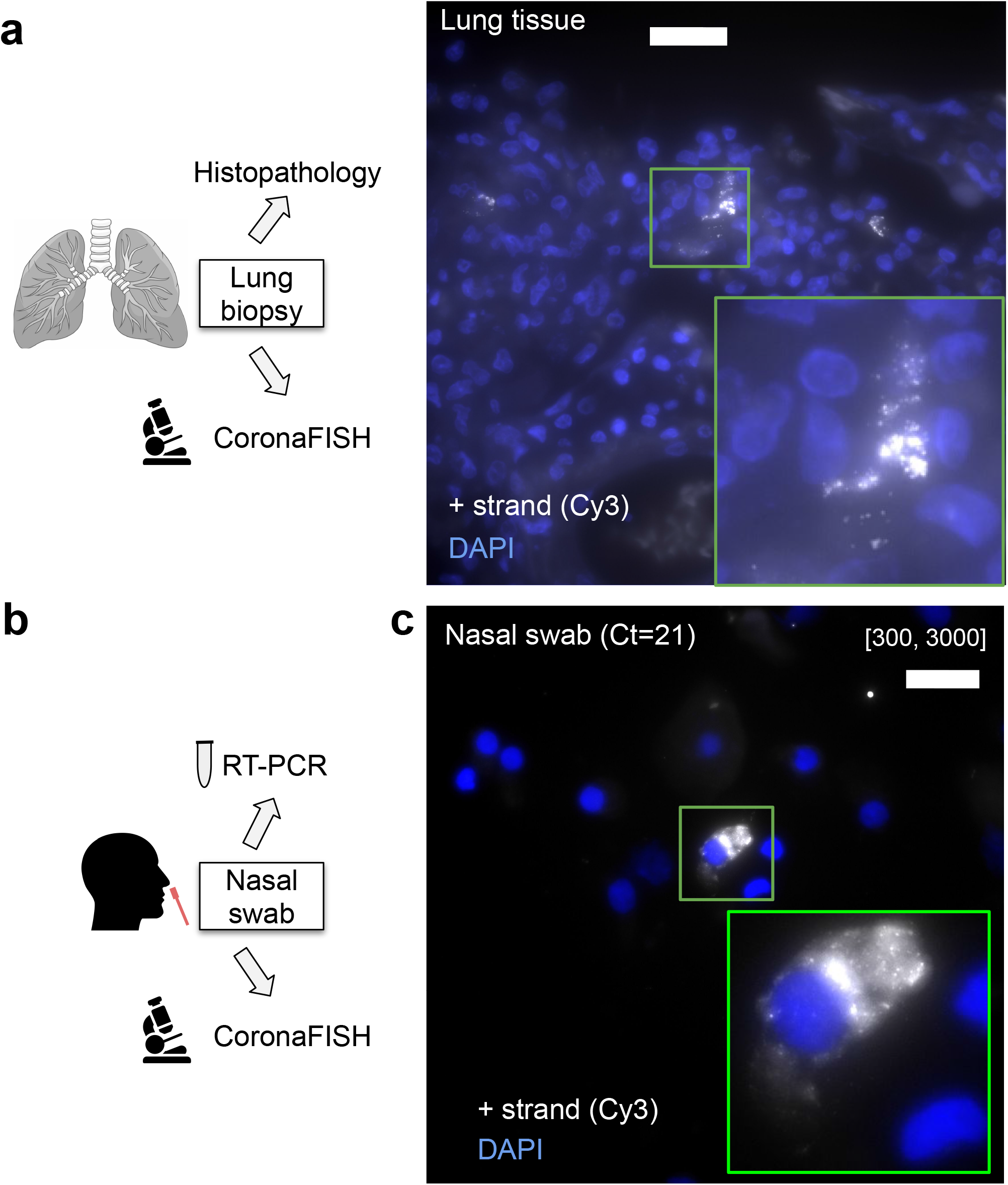
CoronaFISH in human tissue and nasal swabs. **(a)** Detection of positive-strand SARS-CoV-1 in human lung tissue. Scale bar 20 µm. Right image is a magnified view of the green boxed region of interest. Negative control and histopathology images in **Fig EV3. (b)** Nasal swabs were used to perform RT-PCR and the surplus was used for imaging experiments. **(c)** FISH against SARS-CoV-2 RNA in a patient sample tested positive for SARS-CoV-2. Positive-strand RNA was labeled with Cy3 (white), nucleus in blue (DAPI). Scale bar 20 µm. Large field of view and negative controls in **Fig EV2b**,**c**

### Detection of SARS-CoV-2 RNA in nasal swabs

Motivated by the previous results, we next attempted to detect viral presence in human samples used for COVID-19 diagnostics. Nasal swabs were collected from patients with respiratory symptoms as part of routine care at the Hospital Ambroise Paré (Paris) (**Fig 3b**). Samples were screened for the presence of SARS-CoV-2 with RT-PCR. The remainder of the sample not used for diagnostics was deposited on coverglass and we performed CoronaFISH against the positive-strand RNA of SARS-CoV-2. Negative samples (RT-PCR Ct value above detection limit) gave no specific signal, but some areas showed substantially higher background than in the cultured cell lines (**Fig EV3**). This background was rather homogenous, and thus distinct from the RNA signal seen so far in infected cells above. However, in a RT-PCR positive sample (Ct value = 21), we detected a strong RNA-FISH signal in a subset of cells (**Fig 3c**) when staining for the positive strand. Therefore, although a systematic analysis on many more samples will be required to assess specificity and specificity, our CoronaFISH probes may allow the detection of SARS-CoV-2 in patient-derived samples in a clinical setting.

### Electron microscopy visualization of SARS-CoV-2 RNA

Above, we demonstrated how CoronaFISH allows using different fluorescently labeled secondary detector oligos. We reasoned that this flexibility extends beyond conventional fluorescence microscopy, and could also allow for other imaging modalities including electron microscopy (EM). EM is optimally suited to reveal how infection alters the cellular ultrastructure. Indeed, conventional EM images of glutaraldehyde fixed samples showed a dramatic reorganization of the cytoplasm of Vero cells upon SARS-CoV-2 infection, characterized by a loss of Golgi stacks (**Fig EV4a**,**b**) and prominent new features, including numerous DMVs (**Fig EV4c**), which have recently been identified as the main replication organelles of SARS-CoV-2 (Snijder *et al*, 2020; Cortese *et al*, 2020; Klein *et al*, 2020; Wolff *et al*, 2020). Budding of viral particles appeared restricted to the lumen of the endoplasmic reticulum (ER) (data not shown) and to electron-lucent vesicles derived from the ER-Golgi intermediate compartment (ERGIC) that were distinct from DMVs (**Fig EV4d**), also in agreement with prior studies (Sicari *et al*, 2020).

Coupling EM with RNA *in situ* hybridization (EM-ISH) would allow for ultrastructural studies of SARS-CoV-2 infected cells with direct visualization of the viral RNA. We previously used EM-ISH to detect various cellular RNAs using DNA- or ribo-probes (Hubstenberger *et al*, 2017; Yamazaki *et al*, 2018). Here, we adapted this labeling approach by using the same 96 primary oligos against the positive strand of SARS-CoV-2 as before, hybridized to a secondary oligo with Biotin at its 5’ end. These hybrids were detected with an anti-biotin antibody conjugated to 10 nm gold particles (**Fig 4a**). EM imaging was performed on thin sections (80 nm) of Lowicryl K4M-embedded Vero cells, either uninfected or infected (MOI 0.1, 36 h p.i.).

**Figure 4.**
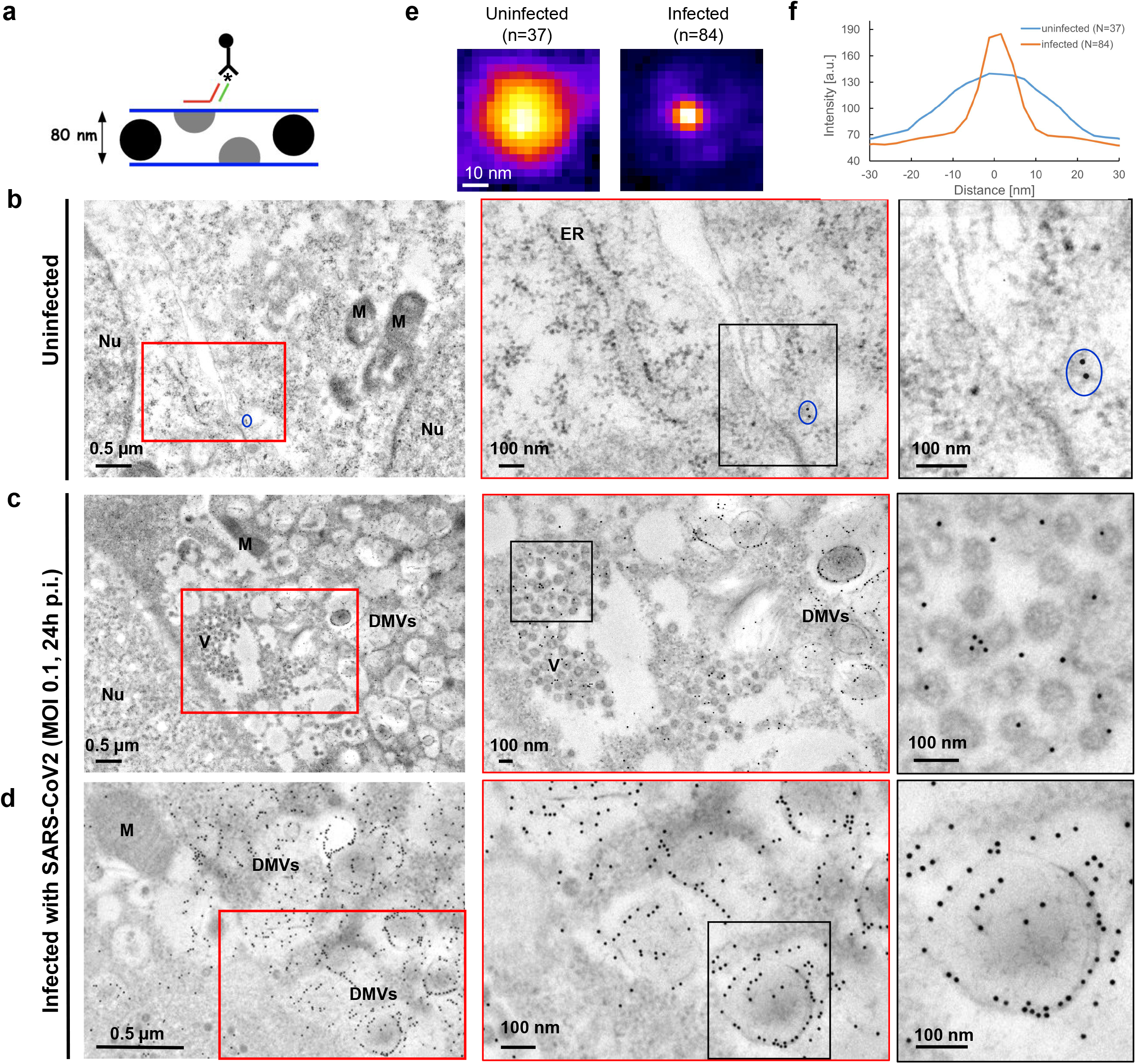
CoronaFISH for electron microscopy. **(a)** Principle of EM-ISH performed on thin sections of Lowicryl K4M-embedded infected and uninfected Vero cells. The asterisk indicates biotin at the 5’-end of the secondary oligo that is recognized by the anti-biotin antibody conjugated to 10 nm gold particles. As sketched, only virions with a section on the upper face of the ∼80 nm ultrathin section will be labelled. **(b)** Uninfected control samples. Blue oval surrounds an example of sparse background staining by electron-dense gold particles. The less defined punctate structures such as those lining up the ER (middle panel) are ribosomes (see panels **e**,**f**). Nu: nucleus; M: mitochondria; ER: endoplasmic reticulum. **(c, d)** Overviews of SARS-CoV-2 infected Vero cells showing major cytoplasmic vacuolization by virus-induced DMVs. Positive strand of SARS-CoV-2 can be detected over intracytoplasmic aggregates of viral particles (**c**). See **Fig EV4e** for extracellular aggregates. DMVs were the most heavily labelled structures, and displayed viral RNAs accumulating especially on peripheral 10 nm fibers (**d**, rightmost frame and **Fig EV4f**). By contrast, mitochondria or nuclei were not significantly labelled. DMV: double membrane vesicles. Nu: nucleus; M: mitochondria. (**e**) Averaged image from punctate structures in panels **b-d** detected with FISH-quant and aligned to the same center (Mueller *et al*, 2013). **(f)** Line profiles through the averaged spots in **e**. The punctate structures visible in infected cells have a size in agreement with the 10 nm nano-gold particles, whereas the punctate structures in uninfected cells are substantially larger and consistent with 30 nm ribosomes.

In uninfected samples, very few gold particles were detected (**Fig 4b**). Manual counting on random fields yielded a mean of only 0.5 +/-0.3 gold particles/μm^2^ in nuclear and cytoplasmic areas (n=9 regions, for a total of 94.4 μm^2^), indicative of low background labeling. Although a much larger number of small particles was visible, their size was consistent with ribosomes rather than gold particles (**Fig 4e**,**f**). In infected cells, gold particles were visible in large numbers at several locations, notably at intracytoplasmic (**Fig 4c**) and extracellular viral particles (**Fig EV4e**), as expected. However, DMVs were the most heavily labelled structures (**Fig 4d**), with manual counts of 180 +/-39 gold particles/μm2 in DMV zones, a 360 fold increase over uninfected cells (n=11 regions, for a total of 11.9 μm2). Strikingly, we observed gold particles accumulating along DMV internal 10 nm thick fibers and at the periphery of DMVs (**Fig 4d, Fig EV4f**). Although the nature of these fibers remains to be determined, this accumulation of gold particles could reflect a slow export of the viral genomes through the recently described pores spanning the DMV double membrane(Wolff *et al*, 2020). Finally, gold particles also labeled large lysosomal organelles, shown to play a role in exocytosis of mouse β-Coronaviruses (Ghosh *et al*, 2020) and containing densely-packed SARS-CoV-2 virions (**Fig EV4f**).

These data demonstrate the flexibility of our probe sets, permitting their use for both fluorescence and EM imaging, and their potential for ultrastructural studies of SARS-CoV-2 replication kinetics.

## Discussion

Here, we presented CoronaFISH, an approach based on FISH (Tsanov *et al*, 2016) permitting the detection of the positive and negative RNA strands of SARS-CoV-2. We validated sensitive and specific detection of SARS-CoV-2 RNA by fluorescence microscopy in Vero cells, several human cell lines (Caco-2, Huh7, and Calu-3), human lung tissue and nasal swabs. Lastly, we demonstrated the flexibility of our approach, by adapting it for EM imaging of the viral RNA.

Our two probe sets each consist of 96 probes, each conjugated to two fluorescent dyes, theoretically enabling 192 fluorescent dyes to target each RNA strand spaced along the entire SARS-CoV-2 RNA. This results in a very bright signal, allowing viral RNA detection in challenging samples, as demonstrated with the patient samples, where the increased signal intensity helps to distinguish true signal despite high autofluorescence. Because our probes span the entire length of the ∼30 Kb viral RNA, they should enjoy high robustness against mutations or partial RNA degradation. Probes are provided individually in a 96-well plate format, therefore subsets of probes against specific regions (e.g. to target specific viral genes) can be selected individually. Further, the secondary detector probes can easily be swapped, allowing the use of different fluorophores and simultaneous imaging of positive and negative strands or even change the imaging modality, as demonstrated by our EM-ISH experiments. This labeling flexibility, together with the fact that our probes target the entire genome (as opposed to only the Spike gene), make CoronaFISH a useful alternative to commercial FISH-based methods for SARS-Cov-2 RNA detection such as HuluFISH and RNAscope.

Compared to immunofluorescence, our hybridization based technique offers several advantages. First, CoronaFISH directly visualizes the viral genome (and/or its replication intermediate) rather than viral proteins. This provides a more specific indication of viral presence and replication, since viral proteins may be found in cells or subcellular compartments where the viral genome is absent and/or where it does not replicate. Thus, CoronaFISH could be instrumental in distinguishing productive from non productive (abortive) infection (Fehr & Perlman, 2015), as has been reported for example in the context of antibody dependent enhancement of SARS-CoV-2 infection (Lee *et al*, 2020). Thereby, CoronaFISH offers a powerful tool to help define the molecular mechanisms of SARS-CoV-2 pathogenesis. In addition, the ability to distinguish and simultaneously visualize positive and negative RNA strands permits the study of replication kinetics in single cells and to uncover spatiotemporal aspects of the infection cycle. Second, the high specificity of these probes owing to their unique complementarity to the SARS-CoV-2 sequence allows to distinguish it from other related viruses, which can be a common problem for antibodies against similar epitopes of different related viral strains. Third, probes can be synthesized within a few days, permitting quick turnover compared to antibody production. Fourth, the CoronaFISH approach is inexpensive. Primary probes can be ordered at low cost, and the provided quantities (smallest synthesis scale provides nanomoles for each oligo) suffice for thousands of experiments. This makes CoronaFISH attractive for high throughput image-based screening of large libraries of antiviral compounds, as illustrated by our Remdesivir experiment.

CoronaFISH can also be used in combination with immunofluorescence for the detection of viral or host proteins (Rensen *et al*, 2020) and is compatible with GFP stains (Tsanov *et al*, 2016). Furthermore, RNA-FISH and IF combined have also been shown to be suitable for flow cytometry and fluorescence activated cell sorting (FACS) (Arrigucci *et al*, 2017). More complex implementations enable multiplexed detection of multiple RNA species (Pichon *et al*, 2018), and will thus permit to probe the host-pathogen interaction at the single-cell-single-virus level.

Compared to the analysis of viral RNA using (single-cell) RNA-seq, CoronaFISH provides information on single cells in their spatial context, since experiments do not require cell dissociation. Our approach can thus deliver insights into the viral life-cycle, including occurrence and abundance of positive and negative-strand RNA, their subcellular localization, and their interplay with the host, as well as with structures induced by SARS-CoV-2 infection (DMVs, replication compartments, ERGIC). CoronaFISH also provides a unique tool to study virus-host interactions in tissue. Furthermore, studying viral RNA presence in thousands of cells is possible by using automated image analysis, and can hence allow the detection of rare events. This will allow examining the effects of infection on aspects such as cell morphology, cell fusion, cell-to-cell transmission, or tissue (re)organization.

Our data on nasal swabs suggest that CoronaFISH may be used on clinical samples and potentially as the basis for diagnostic tests. Unlike standard RT-PCR tests, CoronaFISH does not require RNA extraction or enzymatic reagents, which have at times been in short supply. As mentioned above, because we chose probes targeting the whole length of the SARS-CoV-2 genomes, CoronaFISH detection is also likely to be more robust to mutations, such as the recently identified VUI-202012/01 (aka B.1.1.7) variant, which has been reported to yield negative results in some PCR tests based on the Spike gene (Investigation of novel SARS-COV-2 variant Variant of Concern 202012/01, 2). The cost of CoronaFISH reagents also compares favorably to those used in standard PCR assays. Despite these advantages, a diagnostic test based on CoronaFISH would face two hurdles: speed and a need for a fluorescence microscope. The duration of the FISH experiment (∼2 days) is currently too long for a rapid test. However, microfluidic devices can be used to reduce this delay to less than 15 minutes comparable to fast antigenic tests (Shaffer *et al*, 2015). The requirement for a fluorescence microscope may also be alleviated using cheap do-it-yourself imaging systems, for example smartphones combined with inkjet-printed lenses (Sung *et al*, 2017; Cybulski *et al*, 2014). Such portable and low-cost imaging systems could potentially facilitate point-of-care diagnostics.

We believe that the probes and complementary labeling approaches described here expand the toolbox for studying SARS-CoV-2 and hope that the resources provided (sequences, protocol, and source code) will facilitate their adoption by the community to better understand, diagnose and fight this virus.

## Supporting information

Extra View Figures 1-4

## Acknowledgments

We would like to thank Edouard Bertrand, who originally developed smiFISH, and Hans Johansson for insightful discussions. We also thank Felix Rey for having helped set up the collaboration and Guillaume Dumenil (Ultrastructural Bioimaging UTechS of Institut Pasteur) for having suggested the application of CoronaFISH to electron microscopy and for follow-up discussions. We thank Nathalie Jolly and Nathalie Clément (Center for Translational Science, Institut Pasteur) for their help in obtaining authorization to use patient samples. ESL acknowledges funding from the French Government’s Investissement d’Avenir program, ‘INCEPTION’ (ANR-16-CONV-0005). GBS acknowledges funding by the Institut Pasteur Coronavirus task force (don AXA COVID-19 project COVID-SPREAD). This work was funded by Institut Pasteur.

## Materials and Methods

### Probe design

Entire code for probe-design is available on GitHub: https://github.com/muellerflorian/corona-fish.

The probe-design involves several steps to ensure high sensitivity for the detection of SAR-CoV-2 RNA, while minimizing false positive detection of other β-coronaviruses, other pathogens provoking similar symptoms, and transcripts of the host organism. In parenthesis, we list how many probes remain after each selection step for +strand and -strand targeting probes.

The initial list of probes for the plus and minus strand was generated with Oligostan (N=615 / 608) (Tsanov *et al*, 2016). We then selected all probes with a GC content between 40-60%, probes satisfying at least 2 out of 5 previously established criteria for efficient oligo design (Xu *et al*, 2009) (N=385 / 362).

To guarantee that the probes are insensitive towards known mutations of SARS-CoV-2, we selected only probes with not more than 2 mismatches with any of 2500 aligned SARS-CoV-2 sequences (N=333 / 311).

We then performed a local blast against other β-coronaviruses (SARS, MERS, HKU1, OC43, NL63, or 229E), other pathogens and viruses causing similar symptoms (Mycobacterium tuberculosis, Human parainfluenza virus type 1-4, Respiratory syncytial virus, Human metapneumovirus, Mycoplasma pneumoniae, Chlamydophila pneumoniae, Influenza A-D, Rhinovirus/enterovirus), as well as the transcriptome of the most common host organisms (homo sapiens, mus musculus, African Green monkey, hamsters, ferret). We excluded all probes with more than 22 matches in any of these blast searches (N=115 / 114).

Lastly, we selected the 96 probes with the highest NGS coverage. Probe sequences are provided in **Supplementary Table 1**.

### smiFISH

To visualize vRNA molecules, we used the smiFISH approach (Tsanov *et al*, 2016). Unlabelled primary probes are designed to target the RNA of interest, and can be pre-hybridized with fluorescently labeled secondary detector oligonucleotides for visualization. Probes were designed as described above.

A detailed protocol is available in **Supplementary Note 1**. Cells were fixed in 4% PFA for 30 min, washed twice with PBS++ and stored in nuclease-free 70% ethanol at −20 °C until labeling. On the day of the labeling, the samples were brought to room temperature, washed twice with wash buffer A (2x SSC in nuclease-free water) for 5 min, followed by two washing steps with washing buffer B (2X SSC and 10% formamide in nuclease-free water) for 5 min. The target-specific primary probes were pre-hybridized with the fluorescently labeled secondary probes via a complementary binding readout sequence. The reaction mixture contained primary probes at a final concentration of 40 pm, and secondary probes at a final concentration of 50 pm in 1X NEBuffer buffer (New England Biolabs). Pre-hybridization was performed in a PCR machine with the following cycles: 85 °C for 3 min, followed by heating to 65 °C for 3 min, and a further 5 min heating at 25 °C. 2 µL of this FISH-probe stock solution was added to 100 µL of hybridization buffer (10% (w/v) dextran, 10% formamide, 2X SSC in nuclease-free water). Samples were placed on Parafilm in a humidified chamber on 100 μL of hybridization solution, sealed with Parafilm, and incubated overnight at 37°C. The next day, cells were washed in the dark at 37°C without shaking for >30min twice with pre-warmed washing buffer B. Sample were washed once with PBS for 5 min, stained with DAPI in PBS (1:10000) for 5 min, and washed again in PBS for 5 min. Samples were mounted in ProLong Gold antifade mounting medium.

### Infection of cell lines

The viral stock used originates from BetaCoV/France/IDF0372/2020 and was kindly gifted by the National Reference Centre for Respiratory Viruses at Institut Pasteur, Paris, and originally supplied through the European Virus Archive goes Global platform.

### Vero cells

The day before the infection, Vero cells were trypsinized and diluted in DMEM – Glutamax 10% FBS. They were then seeded, 8*104 / well, in a 12-multiwell with coverslips. The day of the infection the medium of the cells was discarded and the monolayers were infected with SARS-CoV-2 virus in DMEM – Glutamax 2% FBS for 1h at 37°C 5% CO_2_ at a multiplicity of infection (MOI) of 0.02. The MOI used were 0.02. After the desired infection duration, the supernatant was collected for virus titration and the cells were washed with PBS++, fixed with 4% EM-grade PFA for 30 minutes at RT and processed for smiFISH.

### Vero cells for EM-ISH

The day before the infection, Vero cells were trypsinized and diluted in DMEM – Glutamax 10% FBS. They were then seeded, 7 × 10^5^ cells / well, in a 6 multiwell plate. The day of the infection the medium of the cells was discarded and cells were infected with SARS-CoV-2 virus in DMEM – Glutamax 2% FBS at multiplicity of infection (MOI) of 0.1. After 36h, the supernatant was collected for virus titration and the cells were washed with PBS++. The monolayers were fixed using 4% EM-grade PFA in 0.1M Sorensen’s buffer for 1 hour at 4°C.

### Human cell lines

The day before the infection, Huh7, CaCo-2, Calu-3 and Vero cells were trypsinized and diluted in DMEM – Glutamax 10% FBS (Huh7 and Vero), MEM 20% FBS + NEAA, Sodium Pyruvate and Glutamax (CaCo-2), RPMI 20% FBS + NEAA (Calu-3). After 6 hours the medium of the cells was discarded and the monolayers were infected in triplicate with a multiplicity of infection (MOI) of 0.2 with SARS-CoV-2 virus in DMEM – Glutamax 2% FBS. After 36h the supernatant was collected for virus titration and the cells were washed with PBS++. The monolayers were fixed using 4% EM-grade PFA for 30 minutes at RT and processed for smiFISH.

### Titration protocol (Focus Forming Assay)

Vero cells were seeded in 96-multi wells at 2 × 10^4^ cells/well in DMEM – Glutamax 10% FBS. The following day the supernatants to be titered were thawed and serially diluted in tenfold steps in DMEM – Glutamax 1% FBS. 100 ul of the dilutions were used to infect the Vero monolayers, and incubated for 2 h at 37°C 5% CO2. The infection medium was then discarded and a semi-solid media containing MEM 1X, 1.5% CMC, 10% FBS was added to the monolayers. The cells were incubated at 37°C 5% CO_2_ for 36 hours. Cells were then fixed with 4% formaldehyde and foci were revealed using a rabbit anti-SARS-CoV N antibody and matching secondary HRP-conjugated secondary antibodies. Foci were visualized by diaminobenzidine (DAB) staining and counted using an Immunospot S6 Analyser (Cellular Technology Limited CTL). Viral titers were expressed as focus forming units (FFU)/ml.

### Inhibitors assay

To determine IC50 of Remdesivir in our system, Vero cells were pre-treated with serial dilution of Remdesivir (100nM-100µM) for 1h at 37C before infection with SARS-CoV-2 at MOI 0.02. After 2hs infection the virus inoculum was removed, cells were replenished with drug-containing media and incubated for two days. Supernatant was then collected and titered by focus forming assay. IC50 values were calculated by non linear regression analysis (log(inhibitor) vs response – Variable slope (four parameters)) using Prism 6, GraphPad software. For the FISH experiment Vero cells infected at MOI 0.1 were treated with Remdesivir 10 µM and fixed 24hs p.i.

### Image-analysis of infected cell lines

Nuclei were automatically segmented in 2D images with an ImJoy (Ouyang *et al*, 2019) plugin using the CellPose model (Stringer *et al*, 2020). Source code for segmentation is available here: https://github.com/fish-quant/segmentation. Equidistant regions with a width of 50 pixels were calculated around each nucleus. Overlapping regions from different nuclei were removed. 3D FISH images were transformed into 2D images with a maximum intensity projection along z. Signal intensity for each cell was determined as the 90% quantile of all pixels in the equidistant region around its nucleus.

### Lung tissue

Lung autopsy material from the COVID-19 patient was provided by the human biological sample bank of the Lille COVID working group “LICORN” (Lille, France). The use of this autopsy material for research purposes was approved by local ethics review committees at Lille Hospital (Lille, France). Lung autopsy material from the control patient with diffuse alveolar damage prior COVID-19 pandemic was provided by the Pathological department of Necker-Enfants maladies Hospital (Paris, France).

Lung autopsy material was fixed in 10% neutral buffered formalin and paraffin embedded, 4 µm sections were stained with haematoxylin and eosin staining for histological analysis using light microscopy. SARS-CoV-2 RNA was detected as described above.

### Nasal swab patient samples

Respiratory specimens (nasal swabs) have been collected from patients with respiratory symptoms as part of routine care at the Hospital Ambroise Paré (Paris) in late March 2020. No additional samples were collected in the course of this work. Patients were contacted, informed about the research project, and given the possibility to oppose the use of their samples for this project. Lack of opposition to participate in clinical research was verified in the records of all patients whose samples were used here. This project has been recorded in the French public register Health-data-Hub (n°F20200717122429). Processing of personal data for this study complies with the requirements of the “reference methodology MR-004” established by the French Data Protection Authority (CNIL) regarding data processing in health research.

Samples were screened for the presence of SARS-CoV-2 with RT-PCR as described below. Thin-layer preparation from respiratory specimens was achieved through the cytocentrifugation (800rpm, 10min) of 150µL of the remaining respiratory sample with a Thermo Scientific Cytospin 4 cytocentrifuge. Cells were fixed in PBS-PFA 4% for 30 min and then conserved frozen in 100% ethanol until FISH was performed.

### RT-PCR

RNA extraction: 400 μL of clinical samples were extracted in 300 μL of elution buffer (Total NA Lysis/Binding buffer) for 20 min at room temperature with gentle agitation. RNA was extracted with the MagnaPURE compact (Roche) and the MagNA Pure Compact DNA Isolation Kit I (Roche) following the protocol “Total_NA_Plasma_external_lysis purification protocol”. Final dilution of RNA was in 50 µL elution buffer.

RT-PCR: Screening for SARS-CoV-2 was performed by RT-PCR following a modified protocol recommended by the French National Reference Center for Respiratory Viruses, Institut Pasteur, Paris using Ag Path-ID One-Step RT-PCR kit® (Thermofisher). PCR reaction was run on the ABI PRISM® 7900 system (Applied Biosystems) with the following cycle settings: 45°C 10’; 95°C 5’; 45 X (95°C 15’’; 58°C 45’’). Primer sequences and concentration are provided in Supplementary Table 2.

### FISH

Thin-layered samples on a cover-slide suitable for FISH were obtained with a Cytospin protocol. 150 µL of the sample were deposited in a Cytofunnel (Thermo Scientific 1102548). Samples were then centrifuged (800 rpm, 10 min, room temperature) with a Cytospin 4 cytocentrifuge (Thermo Scientific) on Cytoslides (Thermo Scientific 5991059). Cells were fixed in PBS-PFA 4% for 30 min and then conserved frozen in 70% ethanol at −20°C. FISH protocol was performed with Cy3-labeled plus-strand probes labeled with as described above, with one exception: for hybridization 400 µL hybridization buffer was used per sample instead of 100 µL (with the same final probe concentration).

### EM-ISH

#### Fixation and embedding for electron microscopy

For Epon embedding, cell cultures were fixed for 1 hour at 4°C in 2% glutaraldehyde (Taab Laboratory Equipment, Reading, UK) in 0.1 M phosphate buffer, pH 7.3. During fixation, cells were scraped off from the plastic substratum and centrifuged at 5000g for 15 min. Cell pellets were dehydrated in increasing concentrations of ethanol and embedded in Epon. Polymerization was carried out for 48 hours at 64°C. Ultrathin sections were collected on Formvar-carbon-coated copper grids (200 mesh) and stained briefly with standard uranyl-acetate and lead-citrate solutions.

Embedding in Lowicryl K4M (Chemische Werke Lowi, Waldkraiburg, Germany) was carried out on Vero cells fixed either in 4% formaldehyde (Merck, Darmstadt, Germany) or in 2% glutaraldehyde at 4°C. Cell pellets were equilibrated in 30% methanol and deposited in a Leica EM AFS2/FSP automatic reagent handling apparatus (Leica Microsystems). Lowicryl polymerization under UV was for 40 h at – 20°C followed by 40 h at +20°C Ultrathin-sections of Lowicryl-embedded material were collected on Formvar-carbon-coated gold grids (200 mesh) and stored until use.

#### Electron microscopic *in situ* hybridization (EM-ISH)

At the EM level, the SARS-CoV-2 RNA (+) strand probe was composed of the same set of 96 oligodeoxynucleotides that was used for RNA-FISH. The secondary oligonucleotide, however, was modified by a custom-added biotin residue at its 5’-end (Qiagen). Hybrids of the CoV-2 RNAs with the probe were detected with a goat anti-biotin antibody conjugated to 10 nm gold particles (BBI international).

High resolution in situ hybridization was carried out essentially as described previously (Hubstenberger *et al*, 2017; Yamazaki *et al*, 2018). The hybridization solution contained 50% deionized formamide, 10% dextran sulfate, 2 x SSC, and a final concentration of 80 ng/ml of a mix of 1 µg/µl primary oligonucleotides and 1.2 µg/µl of biotinylated secondary oligonucleotide stored at −20°C. EM-grids, with ultra-thin sections of either formaldehyde- or glutaraldehyde-fixed cells side down, were floated for 3 h at 37-42°C on a 1.5 µl drop of hybridization solution deposited on a parafilm in a moist glass chamber. EM-grids were then briefly rinsed over three drops of PBS and incubated 30 min at RT on a drop of goat anti-biotin antibody (BBI International) conjugated to 10 nm gold particles diluted 1/25 in PBS. EM-grids were further rinsed on 2 drops of PBS and finally washed with a brief jet of deionized water at high intensity. Following a 10 min drying on filter paper with thin-sections on top, the EM grids were stained 1 min on a drop of 4% uranyl acetate in water and dried on filter paper 30 min before observation under the EM. Standard lead citrate staining was omitted to favor higher contrast of gold particles over the moderately-stained cellular structures.

Sections were analyzed with a Tecnai Spirit transmission electron microscope (FEI, Hillsboro, OR), and digital images were taken with an SIS MegaviewIII charge-coupled device camera (Olympus, Tokyo, Japan). Quantitation was performed by manually counting gold particles on surfaces that were measured using analySIS software (Olympus Soft Imaging Solutions, Munster, Germany). Statistics were calculated using Excel (Microsoft, Redmond, WA).

