## Supplementary material for "Sensitive visualization of SARS-CoV-2 RNA with CoronaFISH": Extra View Figures 1-4

### for


**Figure EV1: Visualizing SARS-CoV-2 in Vero cells** 2

**Figure EV2: Histology and CoronaFISH imaging of lung tissue** 3

**Figure EV3: CoronaFISH visualization of SARS-CoV-2 in nasal swabs** 4

**Figure EV4: Electron microscopy of infected and uninfected Vero cells** 5

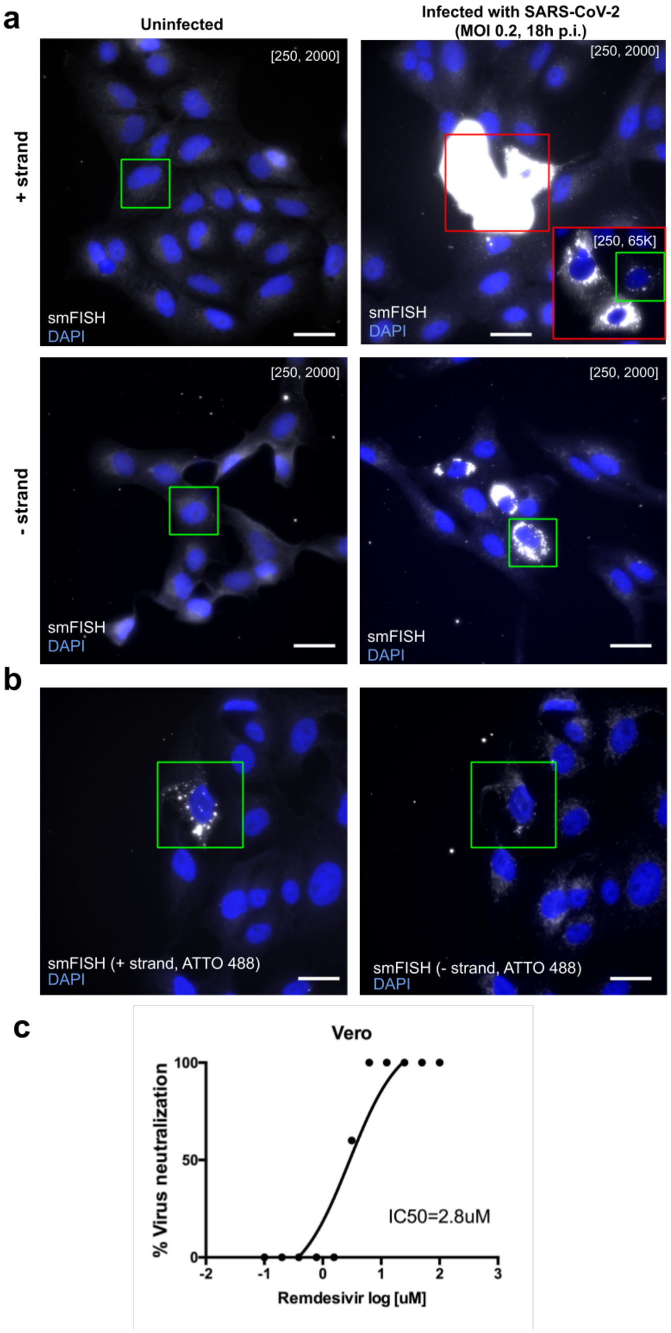


#### Figure EV1: Visualizing SARS-CoV-2 in Vero cells

**(a)** Images of uninfected and infected Vero cells with either the positive or negative strands detected with Cy3-labeled probes. The images show entire fields of view from which insets (green boxes) are shown in **Fig. 1d**. Scale bars 30 μm. **(b)** Dual color imaging of the positive (left) and negative strands (right) of SARS-CoV-2 RNA, as in **Fig. 1f**. Scale bars 30 μm. **(c)** Virus neutralization assay for Vero cells treated by Remdesivir. The calculated IC50 concentration is 2.8 μM.

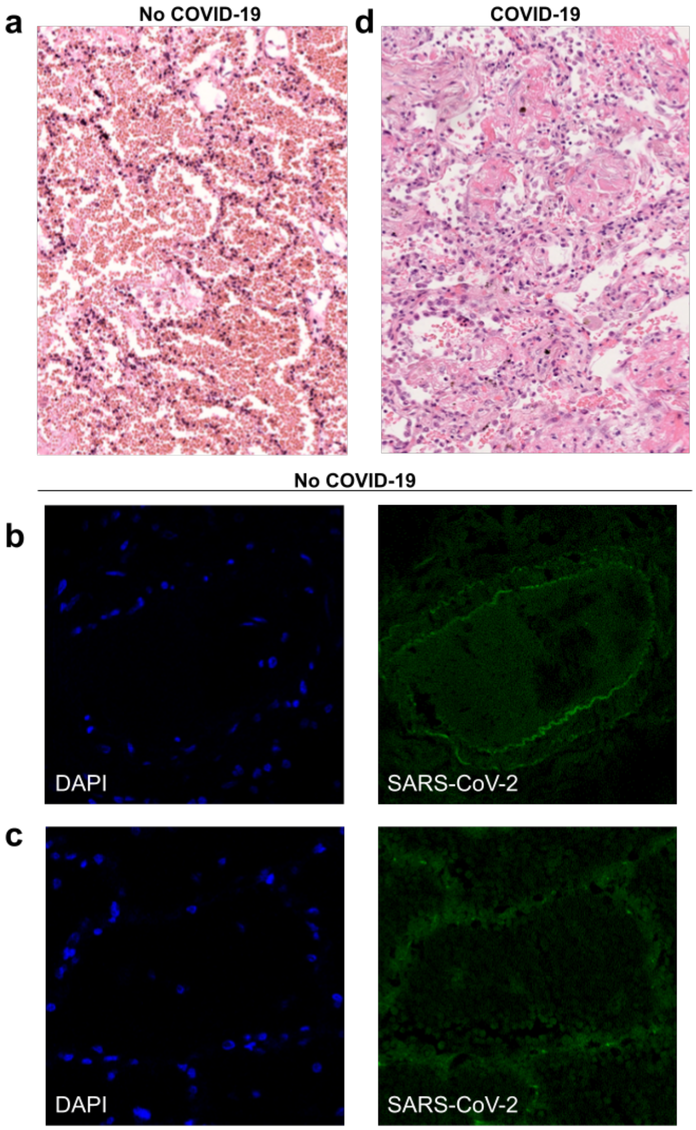


#### Figure EV2: Histology and CoronaFISH imaging of lung tissue

**(a)** Histological analysis of control tissue from a deceased adult patient with acute respiratory distress syndrome (ARDS) obtained prior to the COVID-19 pandemic. Light microscopy (x200) using hematoxylin and eosin staining shows diffuse alveolar damage at exudative phase with an important alveolar hemorrhage, an intra-alveolar and interstitial edema with polymorphic inflammatory infiltrate, and the presence of early hyaline membrane adjacent to alveolar walls. **(b, c)** Confocal microscopy images of a lung arterial section (**b**) and an injured pulmonary alveolus (**c**) of the same patient without COVID-19 as for **a**, with nuclei (DAPI) in blue, and CoronaFISH (positive RNA) probes in green. No intracellular signal is observed with CoronaFISH. The visible extracellular fluorescence is likely due to autofluorescence. **(d)** Histological analysis of COVID-19 lung tissue with diffuse alveolar damage from a deceased adult COVID-19 patient. Light microscopy (x200) using hematoxylin and eosin staining showing diffuse alveolar damage at the organizing phase with intra-alveolar hyaline membranes and fibrin together with interstitial fibrotic lesions with polymorphic inflammatory cell infiltrate of alveolar walls.

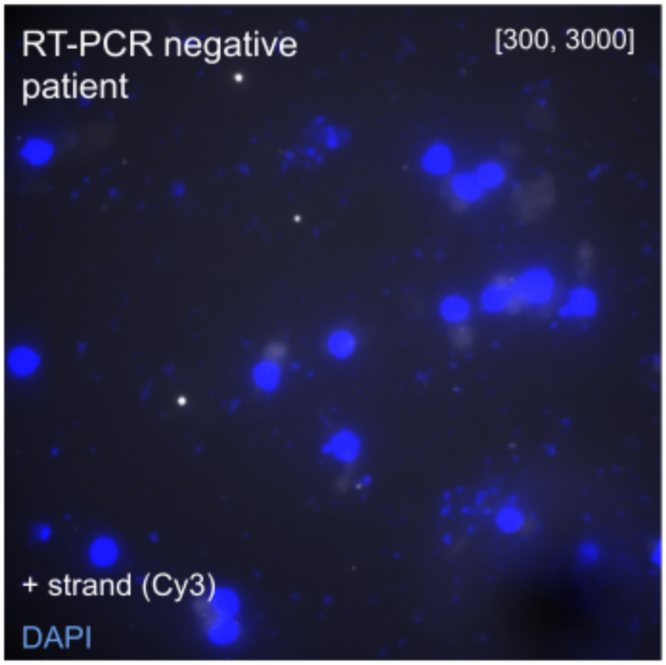


**Figure EV3: CoronaFISH visualization of SARS-CoV-2 in nasal swabs**
FISH against SARS-CoV-2 positive RNA in a patient sample tested negative for SARS-CoV-2 by RT-PCR.

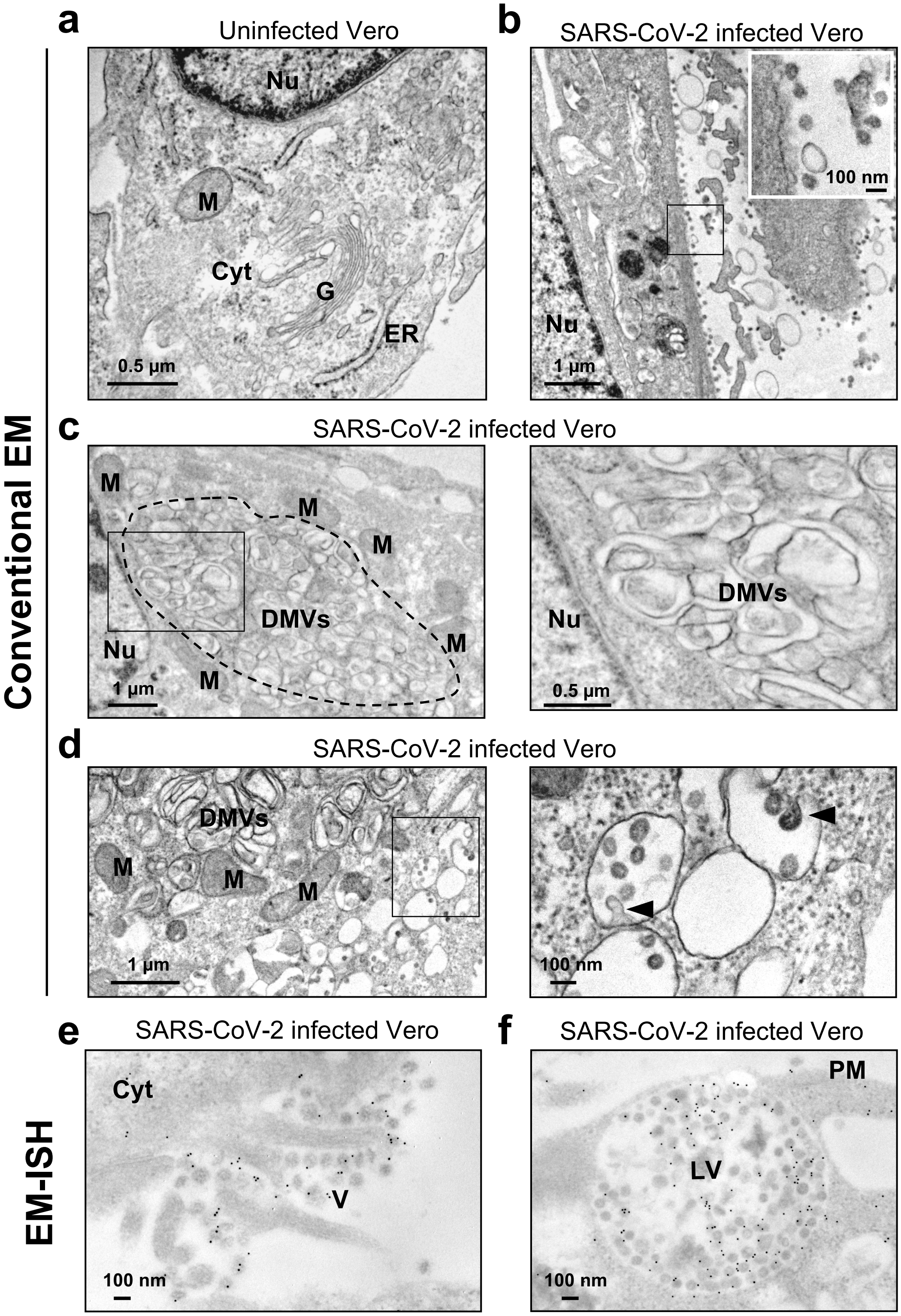


**Figure EV4: Electron microscopy of infected and uninfected Vero cells**
Legend see next page

#### Figure EV4: Electron microscopy of infected and uninfected Vero cells

**(a-d)** Conventional electron microscopy (EM) images (without labeling) of uninfected (**a**) and SARS-CoV-2 infected Vero cells (**b-d**). **b**) Infection leads to a drastic reorganization of the cellular cytoplasm as manifested e.g. by the disappearance of Golgi stacks clearly visible in uninfected cells (**a**) and to numerous viral particles at the plasma membrane (see inset). **c,d**) Infected cells display large regions that accumulate double membrane vesicles (DMVs) (dotted area in **c**), surrounded by mitochondria. **d**) Infected cells also display accumulations of distinct, roundish, electron-lucent vesicles with budding viral particles (arrowheads). **e,f**) EM-ISH images of infected cells with probes against the positive strand of SARS-CoV RNA. **e**) Gold particles labeling the viral RNA can be detected in extracellular virions (right side of the image). **f**) Gold particles labeling densely-packed virions contained within a lysosomal-like organelle implicated in viral egress. Nu: Nucleus; M: mitochondrion; G: Golgi apparatus; ER: endoplasmic reticulum; Cyt: cytoplasm; DMV: double membrane vesicle; PM: plasma membrane; V: Virions; LV: lysosomal-like vacuole.
