## Supplementary Note 1 for "Sensitive visualization of SARS-CoV-2 RNA with CoronaFISH"

### Protocol for CoronaFISH

This protocol is **for tissue cultured cells on cover glass (18 mm) in 12-well plates**. Please adjust buffer quantities to other reaction volumes.


**Reagents** 1
**Buffers** 2
- 4% PFA solution 2
- Hybridization buffer 2
- Wash buffer I 3
- Wash buffer II 3
- DAPI/CellMask staining buffer 3

**Probe preparation** 3
- Primary probes 4
  - Order details 4
  - Prepare probe mix 4
- Secondary probes 4
  - Order details 4
  - Prepare stock solution 4

**Clean coverslips** 5
**DAY 1: fix cells & ethanol permeabilization** 5
**Day 2: hybridization** 5
- **Hybridization solution** 5
- Probe stock solution 5
  - Prepare hybridization solution 6
- Hybridization 6

**Day 3: DAPI staining and mounting** 6
- **Washing and DAPI staining (optional CellMask)** 6
- Mounting 7

**Day 4: Imaging** 7
**Other important aspects** 7

#### Reagents

| Reagents | Reference |
|---|---|
| 32% Paraformaldehyde<br>aqueous solution, methanol free, EM grade | Electron Microscopy Sciences - 15714 |
| Nuclease free water | Ambion™ - AM9922 |
| 10X PBS | Ambion™ - AM9625 |
| 20X SSC | Ambion™ - AM9770 |
| Dextran sulfate | SigmaAldrich - D8906 |
| Formamide | Ambion - AM9342 |
| DAPI | SigmaAldrich - 32670 |
| CellMask™ Green Plasma Membrane Stain | ThermoFisher - C37608 |
| IDT TE pH 8.0 | Integrated DNA Technologies - 11010205 |
| NEBuffer 3 | New England Biolabs - B7003S |
| 70% ethanol molecular grade | SigmaAldrich - 51976 |
| ProLong™ Diamond Antifade Mountant | ThermoFisher - P36970 |

#### Buffers

##### *4% PFA solution*

CAUTION: Work under chemical hood

- 10 ml of 32% stock diluted (1:8)

1. 70 ml 1X PBS (Nuclease free).
2. Add 10 ml PFA 32%.
3. Aliquotes can be stored frozen (@-20°C)

##### *Hybridization buffer*

**CAUTION:** formamide is a teratogen that is easily absorbed through the skin and should be used in a chemical fume hood.

CAUTION: Be sure to let the formamide warm to room temperature before opening the bottle. Aliquot formamide and store at -20°C.

- 100 mg/mL dextran sulfate and 10% formamide in 2X SSC.
- 1 g dextran sulfate (Stored @ 4°C).
- 1 mL 20X saline-sodium citrate (SSC), nuclease-free.
- 1 mL deionized formamide.
- Nuclease-free water to 10 mL final volume.
- Can be aliquoted (e.g. 0.5 mL) and frozen @ -20°C.

1. Mix dextran with 7 mL of water. Vortex and shake O/N to dissolve. Sterile filter.
2. Add formamide and SSC.
3. Add water to final volume of 10 mL.

##### *Wash buffer I*

- MAKE ONLY QUANTITY NEEDED [2 ml per slide].
- 2X SSC

1. 9 mL Nuclease-free water
2. 1 mL 20X SSC

##### *Wash buffer II*

**CAUTION:** formamide is a teratogen that is easily absorbed through the skin and should be used in a chemical fume hood.

CAUTION: Be sure to let the formamide warm to room temperature before opening the bottle.

- MAKE ONLY QUANTITY NEEDED [4ml per slide].
- Can be stored for washes on second day @ 4°C (heat before usage recommended).
- 10% formamide in 2X SSC

1. 20 mL Nuclease-free water
2. 2 mL 20X SSC
3. 2 mL formamide

##### *DAPI/CellMask staining buffer*

- CellMask is optional and only used when cells should be segmented
- CellMask is very bright and can result in bleed through in other channels.

  **Test concentration** carefully before combining with FISH.

1. 10 mL 1X PBS (2 mL for one sample).
2. DAPI 1:1000: 2 µL in 2 mL.
3. CellMask 1:800,000:

a. Dilution 1: 0.5 µL of CellMask stock (2.5µg/µl) to 49.5 µL of PBS.
b. Dilution 2: 1:2000 in PBS, e.g. 0.5 µL in 1 mL PBS.

#### Probe preparation

We recommend using **sterile, DNase/RNase-free low retention reaction tubes**.

##### *Primary probes*

###### Order details

Probes are ordered from IDT in 96 well format. Sequences are specified in an Excel file.

- Plate Product: 25 nmole
- Plate Type: Deep Well
- Standard desalting
- Full yield
- Frozen (on dry ice)
- Concentration 100 µM
- Buffer: IDTE Buffer pH 8.0 (10 mM Tris-HCl/0.1 mM EDTA)

###### Prepare probe mix

1. Primary probes are delivered in 96-well plates. They are provided wet and frozen in Tris-EDTA pH 8.0 (TE) buffer, at final concentration of 100 µM. A provided CSV files contains molecular weight of each probe.
2. Prepare an equimolar mixture of the 24 probes, e.g. 10 ul from each probe mix.
   *Wait until probes reached RT (can take a few hours), gently shake to mix, and spin down on a centrifuge.*
3. Measure probe concentration on NanoDrop. Make 1:10 dilution since stock is too concentrated.
4. Dilute 4-5 times in TE buffer to reach final concentration of 20 *µM*
   *Final concentration of individual probes is 0.833µM.*

##### *Secondary probes*

###### Order details

Secondary oligo for labeling was also ordered from IDT with the following specifications

- Scale: 1 mmole
- Sequence: AA TGC ATG TCG ACG AGG TCC GAG TGT AA
- Modifications: Cy3 on 3', Cy3 on 5'
- Purification: RNAse Free HPLC
- Dry

###### Prepare stock solution

Probes are delivered lyophilized. Resuspend in TE buffer at final concentration of 100 µM. For long-term storage, make aliquots and store at -20°C in the dark.

**NOTE**: Actual amount of probes is less than the ordered scale, please consult the provide specification sheets for the actual amount.

#### Clean coverslips

Different protocols to clean coverslips exist. We use a simple one based on a KOH buffer and a sonicator.

1. Wash coverslips in ethanol, acetone, rinse in water
2. Repeat three times
3. Place the coverslips in 1M KOH 1M in a becher
4. Sonicate during 1h
5. Rinse the coverslip in water until neutral pH; can be kept them in sterile water at 4°C during several months.

#### DAY 1: fix cells & ethanol permeabilization

!!! RNA is sensitive – work with gloves and carefully clean the bench.

!!! Put 70% Ethanol for a few hours @ -20°C

1. Thaw 4% PFA solution, make sure no precipitates are visible (in this case, vortex and heat briefly to 37°C)
2. Wash each sample well with 2 ml of 1X PBS++
3. Remove PBS++
4. Add 1 ml of 4% PFA for each well
5. Incubate 30 min @ RT
6. Wash 2X with 2 ml 1X PBS++
7. Remove PBS++
8. Add 2 ml of 70% ethanol. Make sure coverslip does not float.
9. Seal with Parafilm. Leave O/N @ -20°C.
10. Can be stored @ -20°C for several weeks

#### Day 2: hybridization

##### *Hybridization solution*

!!! Probes are at -20°C; wait until the warm up to RT to avoid condensation, vortex and spin down.

!!! 100 µL of hybridization buffer (containing 2 µL of probe stock solution) is needed for one sample.

!!! We found that only a small excess of secondary probes is sufficient and can help to reduce unwanted background signal.

##### *Probe stock solution*

| **Reagents** | ***Volume*** | ***Final Amount*** |
|---|---|---|
| Probes Primary (20 uM) | 2 ul | 40 pmol |
| Probe FLAP (100 uM) | 0.5 ul | 50 pmol |
| 10X NEB3 | 1 ul | |
| H20 | 6.5 ul | |

| **PCR** [Lid : 99°C] | **Temp** | **Duration** |
|---|---|---|

| | | |
|---|---|---|
| 1 cycle | 85°C | 3 min |
| 1 cycle | 65°C | 3 min |
| 1 cycle | 25°C | 5 min |

##### Prepare hybridization solution

1. Heat hybridization mix to 100°C for 5 min (to dissolve dextran that might not be in solution). Let cool to RT.
2. Add 2 µL of probe stock solution to 100 µL of hybridization buffer, vortex and centrifuge.

##### *Hybridization*

!!! **FORMAMIDE**:

- Don't warm too long. Otherwise it will be ionized. Don't open until it reaches RT.
- Formamide should never be handled without proper safety attire including gloves and goggles.

1. Aspirate the 70% Ethanol
2. Wash twice with 2 mL washing buffer I, and incubate @ RT for 2-5 minutes. Make sure that coverslip doesn't float!
3. Wash once with 2 mL washing buffer II, and incubate @ RT for 2-5 minutes. Make sure that coverslip doesn't float!
4. Create a humidified chamber using a 150 mm tissue culture plate. This chamber will help prevent evaporation of the probe solution from under the cover glass.
5. Place a 10 x10 cm piece of Parafilm in the plate. Press down with forceps and avoid touching.
6. Line the edge of the with a water-saturated paper towel.
7. Within the humidified chamber, dispense 100 µL of the hybridization solution (containing probe) onto the Parafilm. Use forceps to gently transfer the cover glass, cells side down, onto the hybridization solution. Avoid the formation of bubbles.
8. Cover the humidified chamber with the tissue culture lid, and seal it with Parafilm.
9. Incubate @37°C at least for 4h, better O/N, protected from light

#### Day 3: DAPI staining and mounting

##### *Washing and DAPI staining (optional CellMask)*

!!! **Protect** from LIGHT!

1. Heat wash buffer II to 37°C in water bath
2. Add 1 ml washing buffer II to an unused well in the 12 well plate.
3. Transfer coverslip to new 12 well plates with the cells facing UPWARD!
4. Incubate cells in the dark @ 37°C without shaking for >30min.
5. Repeat wash (aspirate buffer and replace with new one).
6. Aspirate washing buffer.
7. Wash 5 min with 2 mL PBS @ RT
8. Incubate 5 min with 2 mL DAPI-CellMask staining buffer shaking.
9. Wash 5 min with 2 mL PBS @ RT

##### *Mounting*

1. Dip slides briefly into water to wash away excess salt
2. Let dry completely (briefly touch the bottom edge to a Kimwipe to remove excess buffer, and then dry them leaning against a rack)
3. Add a small drop (approximately 15ul) of ProLong™ Diamond Antifade Mountant onto a microscope slide.
    - AVOID bubbles. Best is to suck up mounting media by pressing the pipette to the second stop, and release by going only to the first stop.
    - Pipette slowly – it's a little viscous
4. Slowly release coverslip on drop to avoid bubbles.
5. Let harden overnight, protected from light and image cells the day after.
6. Samples can be stored at 4°C in the dark for a few weeks.

#### Day 4: Imaging

Imaging can be performed on a standard widefield microscope with the adequate light source for the used fluorophores. We recommend acquiring 3D stacks to allow for more accurate image analysis.

#### Other important aspects

**Slide handling**

It is advisable to take the habit to always take slides the same way, e.g. with the side with the cells having the same orientation with respect to your forceps. Like this you will always know where the cells are.
